# Sample pooling inflates error rates in between-sample comparisons: an empirical investigation of the statistical properties of count-based data

**DOI:** 10.1101/2022.07.25.501406

**Authors:** Megan N. Taylor, Nic M. Vega

## Abstract

Heterogeneity is ubiquitous across individuals in biological data, and sample batching, a form of biological averaging, inevitably loses information about this heterogeneity. The consequences for inference from biologically averaged data are frequently opaque, particularly when the underlying populations are non-normal. Here we investigate a case where biological averaging is common - count-based measurement of bacterial load in individual *Caenorhabditis elegans* - to empirically determine the consequences of batching. We find that both central measures and measures of variation on individual-based data contain biologically relevant information that is useful for distinguishing between groups, and that batch-based inference readily produces both false positive and false negative results in these comparisons. These results support the use of individual rather than batched samples when possible, illustrate the importance of understanding distributions across individuals within a sample frame, and indicate the need to consider effect size when drawing conclusions from biologically averaged data.

## Introduction

A fundamental concern in data analysis is that of statistical robustness – sensitivity of testing to violation of the underlying assumptions. Robustness is inherently a property of the test applied and the data on which it is used; in particular, robustness of parametric testing against violations of normality in a common concern, as the normal distribution is rarely a good fit to real data (Micceri 1989; Sorrentino 2010; Blanca *et al*. 2013; Mar 2019). Fortunately, many common parametric tests are at least somewhat robust to departures from normality (Rasmussen 1986; Sawilowsky and Blair 1992; Knief and Forstmeier 2021) when other requirements of these tests are met (Erceg-Hurn and Mirosevich 2008), and sturdy non-parametric alternatives are widely implemented (Rasmussen 1986; Potvin and Roff 1993). Further, the central limit theorem (CLT) is commonly invoked to permit the normality assumption for average values. However, the CLT is an asymptotic approximation of the behavior of sample means as the number of measurements becomes large. Formally, the classical CLT does not hold exactly when sampling from a finite population (the finite population CLT instead applies) (Plane and Gordon 1982; Bellhouse 2001), and practically, the definition of “large” depends on the distribution of values within the population being sampled (Smith and Wells 2006; CHANG *et al*. 2008).

Sample pooling or batching, a form of population averaging, is common practice in some areas of biological research. Sample pooling is used to minimize cost and effort in first-pass analysis of population-level signals, for example in population genetics (Dubreuil *et al*. 1999; Lynch *et al*. 2014), vector-based transmission (Ebert *et al*. 2010), and surveillance for contaminants and transmissible pathogens in environmental and agricultural systems (Arnold *et al*. 2011; Bignert *et al*. 2014). In epidemiological surveying, analysis of pathogen load in pooled samples is often used to increase efficiency of screening (Caudill 2010); this approach entered the public eye during the COVID-19 pandemic (Deckert *et al*. 2020). In laboratory experiments, it is sometimes desirable to pool multiple samples when the number of conditions is large and/or the amount of biological material generated per sample is small (Kendziorski *et al*. 2005; Mary-Huard *et al*. 2007). As these conditions are frequently met, batch-based measurements are common. One example is in studies of host-microbe associations in small organisms (Walker *et al*. 2022). In these experiments, “batches” are created by collecting groups of individual hosts for mechanical disruption and quantification, which may include plating on solid agar to determine colony forming unit (CFU) counts. Counts obtained for the batched sample are then divided by the number of hosts in the batch to estimate individual bacterial load. This approach can be used to “average over” heterogeneity in individuals.

Pooling samples for analysis effectively produces a “biological average”. The biological averaging assumption indicates that the value obtained from a pooled sample will represent the arithmetic mean of values in the samples within that pool (Mary-Huard *et al*. 2007). This amounts to an assumption that heterogeneity among samples or individuals is "error" which can and should be smoothed via averaging (Churchill and Oliver 2001; Han *et al*. 2004; Lamichhane *et al*. 2017). This averaging results in loss of information on inter-individual variation within the batch (Bignert *et al*. 1993). When the individual data are normally or log-normally distributed, it is possible to estimate the inter-individual variance from the variance of pooled data (Caudill 2010); however, this is rarely the case for biological data (Micceri 1989).

For these reasons, there has been increasing emphasis on understanding the effects of biological averaging on data. Batching can alter the dimensionality of microbiome communities (Tsilimigras and Fodor 2016), change measurements of diversity (Rodríguez-Ruano *et al*. 2020) and bias measures of total microbial load (Taylor *et al*. 2022a). Further, there is increasing evidence that, even within isogenic and synchronized populations of model organisms, there are biologically significant and replicable patterns of inter-individual heterogeneity (Baeriswyl *et al*. 2009; Gruber *et al*. 2009; Chen *et al*. 2013; Diaz and Viney 2014; Kinser *et al*. 2021), indicating that much of this variation is likely to be information rather than noise or "error".

Here we illustrate the effects of batching using the *C. elegans* host-microbe system as a tractable example, when data are given as counts (colony forming units, or CFUs) per individual. We demonstrate that batching consistently inflates false-positive rates in between-group comparisons, and that inflation of false-negative rates is possible under specific conditions. We investigate the structure of microbial load data for this model host to determine the data properties that allow for inflation of error rates, and we clarify what information is relevant for determining between-group differences in measurements of individual hosts. Based on these results, we provide simple recommendations for experimental design, considering the statistical properties of microbial load data and typical within- and between-group heterogeneity.

### Batching skews inferred counts and inflates false positive rates

Two major issues for batch-based data are: (1) batching is an average (arithmetic mean) over individuals, which inherently involves some loss of information about the distribution of individual observations and (2) it is difficult to interpret the arithmetic mean as a population measure if data are not symmetric and centrally distributed. This can be demonstrated directly by comparing individual colonization data to batch-inferred CFU/worm data from the same experiment. For example, in N2 worms colonized with *Staphylococcus aureus* Newman, batching pulls estimated CFU/worm toward the upper quantiles of the distribution, and this effect becomes more marked as batch size increases (20 individuals/batch vs 5 individuals/batch, **Figure 1A**). This reflects the right-hand skew in the individual data (median<mean; here, median=5,500 and mean=20,884).

**Figure 1.**
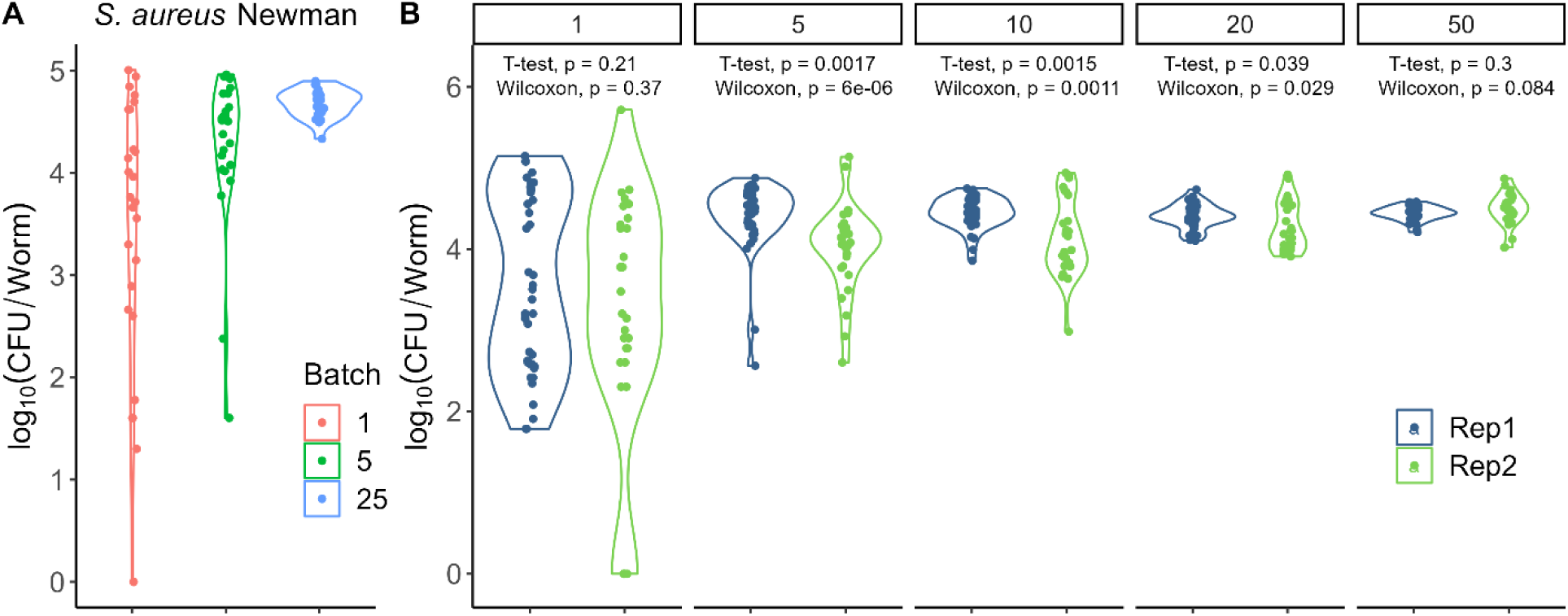
Batching individuals for CFU/worm quantification approximates the sample mean and amplifies differences in run-to-run variation in the distribution of data. (A) Experimental batching skews estimated CFU/worm toward the upper quantiles of the data. Here, synchronized N2 adult hermaphrodites were colonized by feeding on NGM plates with lawns of *Staphylococcus aureus* Newman for 20 hours. Worms were cleaned and permeabilized according to standard protocols (Methods), then mechanically disrupted as individual worms, batches of 5 worms, or batches of 25 worms for dilution plating and CFU/worm quantification (n=24 data points each). Data are CFU/worm, as measured for individuals (batch size 1) or as inferred by dividing the CFU count for each batch digest by the number of individuals in that batch. (B) Simulated batch data, created by resampling individual-worm bacterial load data within runs. Raw colony count data from two independent runs of the same experiment (N2 adults colonized with *Salmonella enterica* LT2, original data shown as batch size 1) were bootstrapped with Poisson error of counts (see Methods) to create 25 simulated data points representing batches of 5, 10, 20, or 50 worms per measurement. Tests indicate the result of comparisons of run 1 vs run 2 data at each batch size.

We assess the practical importance of this effect in simulations, by drawing from single-worm data to create simulated “batch digests”. Here we use two runs of the same experiment (biological replicates, performed on different days; n=48 worms per run; **Figure 1B**). In these experiments, N2 adults were colonized on NGM agar plates with *S. enterica* LT2 for 48 hours before measurement of bacterial load in individual worms. Individual-worm data were not significantly different between runs. In both runs, the data showed a right-hand skew (median < mean), but the mean and skewness were different (run 1 mean 27,497 and median 3,200; run 2 mean 29,207 and median 1,600).

We performed a series of bootstraps on these data (resampling the raw CFU count data with replacement and imposing Poisson error of counts; see Methods) to determine whether batch digests would have an impact on assessment of run-to-run variation. Simulated batches of different sizes (5, 10, 20, or 50 worms per batch, **Figure 1B**) behaved as expected from **Figure 1A**. In accordance with expectations from central limit theorem, as batch size increased, the variance between measurements decreased and the inferred CFU/worm was pulled toward the arithmetic mean of the individual-worm data, which is located toward the upper quantiles of the right-skewed distribution. Further, run to run differences in individual-worm data clearly affected the behavior of the distributions as batch sizes increased. Comparison of CFU/worm across runs in bootstrap-based simulated batch data indicated that differences between runs were interpreted as significant in batched data. The situation was not improved by instead comparing log_10_-transformed counts (t-test batch 5 p=0.02, batch 10 p=0.002, batch 20 p=0.04, batch 50 p=0.3; Wilcoxon tests unchanged). For these data, increasing batch size altered the conclusions of the analyses, indicating with increasing frequency that there was a significant difference between biologically replicate runs of the same experiment.

As in previous work from this lab (Taylor *et al*. 2022a), batch digests introduced spurious differences between runs in the *Salmonella* CFU/worm data (false positive). However, a larger batch size was required to produce elevated false positive rates here (using the *Salmonella* data set) than in the previously published example, which used single-worm colonization data for different bacteria (MYb53 and MYb120 from the *C. elegans* native microbiome (Dirksen *et al*. 2016)). This raised the question of why, and what features of these data are responsible. Understanding the origins of batch-based errors in these data will be important for determining when batching is and is not admissible.

### Quantifying run-to-run heterogeneity

At some magnitude, it is reasonable to consider a shift in the distribution of values as a true difference between groups, whether or not the arithmetic means differ. This raises two questions: First, how much of a change in average and in distribution is expected across replicates of the same experiment or sample type (as any between-group differences smaller than this will not be interesting), and what is the best way to quantify this expectation? Second, how does the distribution of the data alter the error structure of between-group comparisons, and what does this mean for the ability to accurately detect differences?

To address the first question, we re-analyzed existing data to determine the distributional properties of CFU/worm data. A large set of single-worm measurements collected as part of a study on heterogeneity in a minimal native worm microbiome (Taylor and Vega 2021) was filtered to remove all replicates containing less than 18 individuals to ensure sufficient sampling of inter-individual heterogeneity, leaving data representing eight host genotypes with at least two replicate data sets for comparisons. Mono-colonization data for *Salmonella enterica, Staphylococcus aureus, and Microbacterium oxydans* in N2 adults were added to this data set, for a total of 11 experimental conditions with replicates performed on separate days **(Figure 2A**).

**Figure 2.**
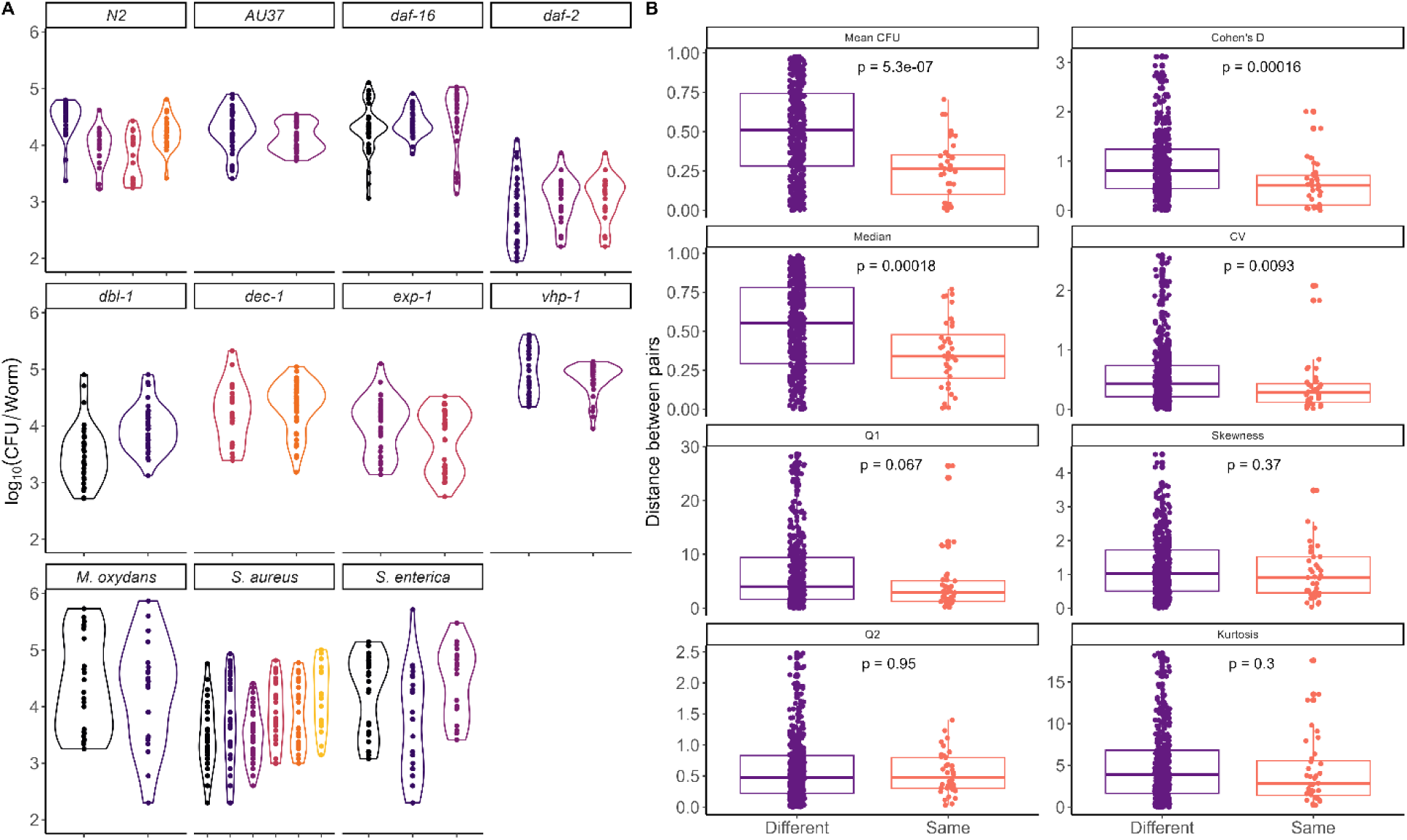
Comparisons of CFU/worm data within and between conditions. (A) CFU/worm data for each condition x replicate (host genotype for minimal community data, first two rows; colonizing bacterial species for mono-colonization data in N2 adults, bottom row). Each point represents an individual worm. Data are colored by replicate. (B) Pairwise differences in summary statistics from (A), grouped according to whether the runs compared are replicates of the same condition (same host genotype for minimal microbiome data or same colonizing microbe for single-species colonization data, "same", in red) or from different conditions (purple). Cohen’s D is (M1-M2)/s, where Mi is the mean of sample 1 or 2 (M1>M2), and s is the smaller of the two standard deviations. Hogg’s statistics are calculated as Q1=(U_0.05_-M_0.5_)/(M_0.5_-L_0.05_) and Q2=(U_0.05_-L_0.05_)/(U_0.5_-L_0.5_), where U_0.05_ and L_0.05_ are the means of the upper and lower 5%, M_0.5_ is the mean of the middle 50%, and U_0.5_ and L_0.5_ are the means of the upper and lower 50% of the data. Q1 is a measure of skewness, and Q2 is a measure of tail weight. Distances between means and medians are the rescaled quantity obtained by dividing the absolute difference between the statistics of two replicates by the sum of the statistics of those two replicates. Distances in the scale-free quantities CV, Q1, Q2, skewness, and kurtosis are the absolute value of the difference across pairs of replicates. Tests shown are Wilcoxon rank sum tests comparing distributions of “different” vs “same” pairs for each summary measure.

Measures of central tendency (mean, median, Cohen’s D) and coefficients of variation for CFU/worm data were more similar run-to-run across replicates of the same experiment (“same”) than across experimental conditions (“different”, **Figure 2B**), suggesting that these statistics contain information useful for distinguishing between groups. This comparison was not significant in the higher moments of the distributions (skewness and kurtosis), although these measures are known to be sensitive to outliers, and Hogg’s Q1, a more robust measure of skew, showed a marginally significant difference (Hogg 1974; Hill and Dixon 1982). These higher moments may therefore be more usefully considered as general properties of this kind of data. Overall, these data tend strongly to be heterogeneous (0.5<CV<1.5), asymmetric (nearly always skewed to the right) and heavy-tailed (positive excess kurtosis), with indications of multi-modality (**Figures S1-S2 and Table S1, File S1**).

### Data heterogeneity and false positives

It is easy to demonstrate (**Figure 1; Figure S3 and Tables S2-S4, File S1**) that batching, as a form of biological averaging, produces data that behave mathematically as an average with all the properties that entails. When two populations have the same mean, even when the distributions of individuals within these populations are very different, batch-based sampling can result in no difference between samples being detected (false negative; **Table S4, File S1**). However, in the real data examples so far, batching tended to produce inflated false positive rates. It is illustrative to look at the properties of data to isolate features that will promote false positives vs. false negatives.

To do so, we employed a set of simple simulations (**SI**) using the flexible beta distribution. First, we considered the case where data sets have the same mean but different distributions (**Figure S3**). This case is useful because we know what commonly-used statistical tests should report for the individual-based data - the null hypothesis of the t-test (no difference in means) is true even when the null hypothesis of the Mann-Whitney test (no difference in the sum of ranks) is not. While sampling and small number effects produced an expected rate of false positives in simulated data with a common mean (**Table S4, File S1**), this was insufficient to produce *elevated* false-positive rates due to batching.

For real data, even under ideal conditions, it is unlikely that means will be exactly the same across biological replicates. Data in replicates will differ slightly due to run-to-run differences (small shifts in timing, pipetting noise, etc). In the *Salmonella* colonization data, for example, average colonization in run 1 was 94.1% of the average colonization in run 2 two weeks later and 53% of that from a third run almost a year later with a different operator (**Figure 2A**, all Mann-Whitney U tests p > 0.05). This is typical of this type of data; normalized distance between average values across replicates showed a high-density region around 0.25 with few comparisons above this value, corresponding to a ∼50% difference when averages are similar (**Figure 2B**).

Allowing small run-to-run variation in simulation parameters (**Figure S4**) produced results consistent with real data. The between-run distances for moments of the individual-"worm" data in these simulations were consistent with observed values, suggesting that these simulations adequately represented relevant features of the data (**Figure 2B**, **Figure 3**). Also consistent with results from real data, in simulations with run-to-run variation, batching promoted false-positive results in pairwise comparisons (**Table 1**). The increased rate of false positives with batching was not due to dispersal of the means in batch data (**Figure S4, File S1**). Nor did batching strongly affect the between-run distance in skew or any properties of the excess kurtosis (**Figure S4, File S1**). However, variation (shown as CV, **Figure S4, File S1**) decreased as batch size increased, consistent with expectations from previous sections and from the central limit theorem. As data points clustered closer to the mean, run-to-run variation in that mean (which in simulations was due to small random variation in the parameters of the underlying distribution) was interpreted as a real difference between groups. Notably, the t-test and the Mann-Whitney test behaved similarly; the parametric test reacts to the collapse of uncertainty about the value of the mean, and the nonparametric test reacts to the coalescence of the distribution of biologically-averaged data points toward the mean. The conclusion of significant differences across runs is in some sense true (small fluctuations in simulation parameters produce means which are not identical) but is not meaningful (the underlying base parameters are unchanged; fluctuations in these parameters are small and would optimally be dismissed as insignificant).

**Figure 3.**
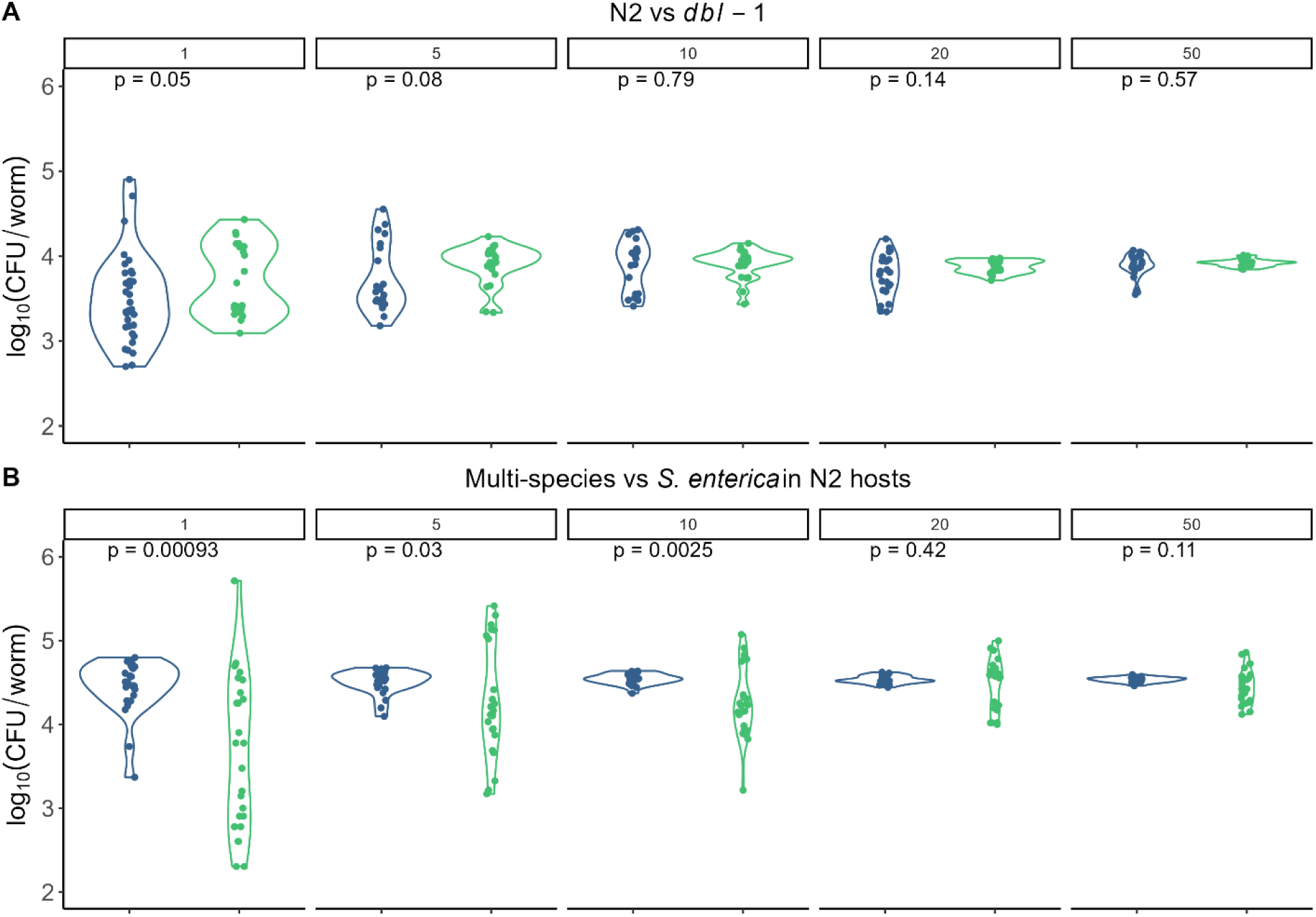
Raw data (shown as batch size 1) and example simulated batch digests (5, 10, 20, and 50 individuals/batch) generated by bootstrapping (A) multispecies total CFU/worm data for N2 (blue) vs. *dbl-1* (green) adults and (B) multispecies total CFU/worm (blue) vs *S. enterica* (SE) total CFU/worm (green) in N2 adults. Raw data sets are each one replicate of a given experiment (n=22-24 worms); number of simulated data points at each batch size is the same as the smaller of the data sets. Tests shown are Wilcoxon rank sum tests comparing each pair of simulated data.

**Table 1.** Results of comparisons of two sets of data drawn from beta distributions with the indicated baseline parameters, where parameter values (α,β) are drawn independently for each simulated data set within each run of simulations (1000 runs, max 10^5^ CFU/worm, 24 data points at each batch size, parameter deviances drawn from U[-0.1, 0.1]). Data shown are fraction of runs where p<0.05 (false positive).

|  |  | Batch Size |  |  |  |  |
| --- | --- | --- | --- | --- | --- | --- |
|  |  | 1 | 5 | 10 | 20 | 50 |
| Beta(1, 5) | t-test | 0.07 | 0.15 | 0.23 | 0.33 | 0.52 |
|  | Wilcoxon | 0.06 | 0.14 | 0.21 | 0.32 | 0.51 |
| Beta(0.5, 2.5) | t-test | 0.08 | 0.09 | 0.14 | 0.22 | 0.40 |
|  | Wilcoxon | 0.07 | 0.09 | 0.12 | 0.23 | 0.39 |
| Beta(5,1) | t-test | 0.08 | 0.13 | 0.20 | 0.32 | 0.50 |
|  | Wilcoxon | 0.07 | 0.13 | 0.20 | 0.31 | 0.49 |
| Beta(2.5, 0.5) | t-test | 0.05 | 0.10 | 0.14 | 0.25 | 0.40 |
|  | Wilcoxon | 0.06 | 0.09 | 0.14 | 0.24 | 0.40 |
| Beta(5, 5) | t-test | 0.10 | 0.25 | 0.40 | 0.53 | 0.69 |
|  | Wilcoxon | 0.10 | 0.25 | 0.39 | 0.51 | 0.68 |
| Beta(2.5, 2.5) | t-test | 0.07 | 0.19 | 0.29 | 0.44 | 0.60 |
|  | Wilcoxon | 0.07 | 0.17 | 0.28 | 0.42 | 0.59 |
| Beta(1, 1) | t-test | 0.07 | 0.11 | 0.18 | 0.30 | 0.47 |
|  | Wilcoxon | 0.06 | 0.10 | 0.18 | 0.29 | 0.46 |

To understand what data properties might affect false positive rates during biological averaging, we considered how the higher moments of the simulated data affected these comparisons. The impact of kurtosis (tail weight) was minimal, as can be seen by comparing the effects of batching when parameters are centered around Beta(1,5) vs. Beta(0.5, 2.5) (**Table 1**). Likewise, the direction of skew had minimal effect, as when the values of α and β were reversed (**Table 1**). However, the magnitudes of both variance and skewness were important, and to some extent it was possible to trade one for another; symmetric distributions were more likely to produce significant differences at a given batch size than skewed distributions with similar variance (compare for example Beta(1,5) with variance 0.02 and skewness 1.18 vs. Beta(5,5) with variance 0.02 and skewness 0, **Table 1**). The reason for this is surprisingly straightforward: for symmetric distributions, the data points are clustered closer to the true within-run mean (which is not precisely identical in simulations due to run-to-run variation in underlying parameters) than for asymmetric distributions. This is true even in individual data but is exacerbated by batching (**Figure S5, File S1**). This result recalls the Berry-Esseen theorem in statistics, which indicates that the rate at which the arithmetic mean converges to a normal distribution is a function of the sample size (as n^-1/2^) as well as the ratio of the third and second absolute normalized moments of the distribution of the random sample (Berry 1941; Esseen 1942; Korolev and Shevtsova 2010).

### Introducing experimental realism: False-positive rates and run-to-run variation in data with replicates

Real experimental data have additional structure. Data are generally combined across multiple replicates, and most experiments are not large enough to create many batches of 20-50 individuals per condition. Simulations were therefore performed to assess the effects of replication on false-positive rates (**Table S5 and Figures S6-S7, File S1**). Briefly, these simulations indicated that experimental designs including replication did not decrease false-positive inflation due to batching.

### Data heterogeneity and false negatives

If false positives can be obtained when heterogeneous data from the same base distribution are biologically averaged (**Figure S4, File S1**), false negative rate inflation can occur when samples have different base distributions but similar means. We concluded earlier that averages were overall more similar across runs within the same type of experiment than across experiments in the example data. However, there is substantial overlap (**Figure 2**), and our example data set includes several instances where data with very different distributions have similar means.

Based on previous results, we expected to get false negative inflation for biologically averaged data if distributions of individuals were differently variable and/or skewed but had sufficiently similar means. An example was provided by two data sets representing host genotypes Bristol N2 (n=22; average 8,277 CFU/worm) and *dbl-1(nk3)* (n=33; average 7,799 CFU/worm). Here the N2 bacterial load data were moderately variable and skewed (CV 0.88, skewness 0.89) as compared with the more heterogeneous *dbl-1* data (CV 2.06, skewness 3.37), and the two data sets were distinguishable from one another (original data shown as batch size 1 in **Figure 3A**). We resampled these data to simulate batching as before (n=22 data points in all simulations to match the smaller of the original data sets, **Figure 3A**). Over 10,000 runs of this resampling, we observed the correct result of p<0.05 in (38.2%, 19.7%, 16.9%, 9.5%) of runs at batch sizes of 5, 10, 20, and 50 worms respectively. As a second example, we compared one run each of *S. enterica* colonization (n=25, mean 33,880, CV 3.03, skewness 4.22) vs. minimal microbiome colonization (n=24, mean 34,069, CV 0.49, skewness –0.07), both in N2 adults. Comparison of the original data sets (shown as batch size 1 in **Figure 3B**) indicated a difference between conditions (Wilcoxon p= 0.0009). When we resampled these data as before (10,000 runs, each run with n=24 data points at each batch size to match the smaller data set, **Figure 3B**), we again saw false-negative rate inflation; true positive p<0.05 was achieved in 93.4%, 51.1%, 10.2%, and 12.2% of runs at batch sizes of 5, 10, 20, and 50 respectively. False negative rates were therefore inflated by batching.

Run-to-run variation in samples can affect false negatives, but this effect is likely to be a matter of happenstance. If the distributions of conditions A and B have similar means but different variance and skew, across any number of runs, false negatives will be a potential issue in batch data when sample averages happen to be sufficiently similar. Importantly, if only biologically averaged data are taken and the underlying distribution is not known, there is no obvious way to determine whether this is likely to have occurred for any given comparison.

## Discussion

Experimentalists are regularly warned by statisticians that if an inappropriate test is used, the conclusions of data analysis may be incorrect (Bridge and Sawilowsky 1999; Othman *et al*. 2004; Vickers 2005; Erceg-Hurn and Mirosevich 2008; Zuur *et al*. 2010; Forstmeier *et al*. 2017). The results shown here are an example of the empiricist’s addendum – if the data are problematic, the test is the lesser problem. Commonly, data can be low-quality due to sampling issues, bias, mislabeling, contamination, etc. (Cortes *et al*. 1995; Clausen and Willis 2021; Schloss) In the case illustrated here, data quality issues are not necessary to produce spurious results. Instead, this case belongs to the family of problems where sampling the wrong thing – or measuring the right thing in the wrong way – produces measures unsuitable for answering the question being asked (Riniolo and Porges 2000).

We use real data and simulations to illustrate a case where error rates are inflated by biological averaging. For both false positives and false negatives, errors arise because batching produces data points representing the arithmetic mean of the individuals included in the batch. As batch size increases, the central limit theorem indicates that the distribution of these data will lose information about the original distribution and become centered around the sample mean. When distributions are similar, as in experimental replicates, run-to-run variation can create variation in sample means that may be mistaken for a real difference. When distributions are different and when the means of two groups happen to be similar, batching-associated loss of information about the distributions of the data can lead to failure to distinguish between groups. Critically, if information about the distribution of individuals is missing and particularly when effect size is small, there is no obvious way to determine whether a given comparison is likely to experience inflated rates of error.

The results shown here illustrate the fact that batching, by producing a biological average, generates values that converge toward the arithmetic mean. This should not be interpreted to mean that these batch-inferred values are distributed as Gaussian – even without testing for normality, it is visually obvious that this is not true for the data used here (**Figures 1, 3**). It is known that convergence to normality as sample size increases is slow for variable and skewed data (Korolev and Shevtsova 2010). Further, empirical work has demonstrated that discreteness of data and skewness and kurtosis of a distribution dramatically increase the sample size required for the population of means to approach normality (Smith and Wells 2006; CHANG *et al*. 2008). For log-scale, skewed, and heavy-tailed data typical of bacterial load measurements, even at large batch sizes the distribution of the population of means is clearly contaminated (**Figure 3**), with marked asymmetry and even multimodality. We do not attempt to describe this progression formally. For the purposes of the present analysis, it is sufficient to note that the population of means loses biologically relevant information about the underlying heterogeneity among individuals, while failing to gain the advantages conferred when normality can be assumed.

As part of this analysis, we describe common features of CFU/host data in the small model host *C. elegans* and confirm measures that are useful for distinguishing between groups. The lower moments of the data (mean and variance) and the central quantiles are most strongly associated with specific combinations of host strain and microbial colonist(s), suggesting that both central measures and measures of variation contain information that is useful for distinguishing biologically relevant groups. Further, these data are almost always right-hand skewed and frequently heavy-tailed – these appear to be general features of this data type. However, due to the relatively small *n* within replicates in these data, it is not at present possible to rule out the possibility that specific measures of asymmetry and/or tail weight might also contain distinguishing information in larger samples. Hogg’s Q1 and Q2, which were designed as stable measures of skewness and tail weight, are promising for sufficiently large samples where the 5% and 95% quantiles can be repeatably described. Interestingly, single-species colonization by the two pathogens in this data set is associated with higher CV and broader range than colonization by members of a native microbiome, as well as with a more pronounced multi-modality; additional data, including information on single-species colonization by commensals, will be required to determine generality of this observation.

For the experimentalist, these results provide guidelines to minimize the risk of false-negative and false-positive comparisons. Heterogeneity between individuals (here, observed even within populations of isogenic, synchronized worms) means that individual-based data should be preferred over batched data if possible. Effect size should be considered along with statistical significance when making comparisons between groups (Sullivan and Feinn 2012), as small but statistically significant differences are likely to be enriched in false-positive comparisons, particularly if batching cannot be avoided. The underlying distribution of the data (variation and centrality/skewness in particular) and the run-to-run variation will be important – the results shown here (**Figure 2**) indicated typical run-to-run variation of around 50% in average CFU/worm (normalized distance of means <0.25, or Cohen’s *d* <1.3), suggesting that run-to-run effects should be estimated for a given system and used to determine thresholds for effect size.

More broadly, these results suggest a need for the experimentalist to consider carefully the question that is being asked. First, is the question about what happens on average? Batching by its nature averages across individuals. If the question is not about the population average (arithmetic mean), batch-based data are perhaps not best suited to provide an answer. There are many cases where the arithmetic mean is not the most informative quantity to describe a process. For example, “bet-hedging” in a fluctuating environment is canonically represented by geometric mean fitness (Starrfelt and Kokko 2012), and transmission of infectious disease is often driven by “superspreading” with a long-tailed distribution of secondary infections (Woolhouse *et al*. 1997; Galvani and May 2005; Lloyd-Smith *et al*. 2005; Paull *et al*. 2012; VanderWaal and Ezenwa 2016). Further, instability of the arithmetic mean in asymmetric and “lumpy” data has been historically described (Ansell 1973; Hill and Dixon 1982).

Second, is it acceptable not to observe individual heterogeneity? Even if we assume the average is the correct summary statistic for a given question, variation may still be relevant. We observe (**Figure 2**) that heterogeneity in bacterial load has characteristic features that are repeatable across replicates but change across experimental treatments, inconsistent with the idea that this heterogeneity is simply attributable to sampling noise. As stated above, measures of variation among individuals appear to be characteristic, indicating that sacrificing individual heterogeneity can lead to loss of information that is useful for distinguishing between groups.

It may be useful to extend these analyses to other common data types with different structures. In *C. elegans*, count data are commonly taken in studies of fecundity (offspring per hermaphrodite), and mortality data are taken as counts and presented as fractional survival over time. Both data types are well known to have issues related to hidden heterogeneity within populations (Wu *et al*. 2006; Suda *et al*. 2009; Baeriswyl *et al*. 2009; Gruber *et al*. 2009; Pincus *et al*. 2011; Chen *et al*. 2013; Diaz and Viney 2014; Kinser *et al*. 2021; Travers *et al*.). These data also have characteristic complications, notably censoring, which have well-known relevance for analysis (Escobar and Meeker 1992; Gijbels 2010) but which are minimal in the bacterial load data used here. If (as seems probable) variation in these data also contains information about the underlying biology, understanding inter-individual and inter-sample heterogeneity will be important not only statistically, to allow appropriate choice and interpretation of analyses, but also conceptually, to allow experimental design that captures this information and conclusions that take heterogeneity into account. Understanding the nature of heterogeneity in data therefore presents an opportunity to maximize the potential of an experimental model by improving accuracy and repeatability of results.

## Acknowledgements

This research was funded by Emory University and NSF [grant number PHY2014173]. The authors would like to thank C. LaRock for generously providing us with bacterial strains, and L. Waller and L. Morran for discussions and feedback on this manuscript.

## Methods

### Strains and materials

*C. elegans* strains were obtained from the *Caenorhabditis* Genetic Center, which is funded by NIH Office of Research Infrastructure Programs (P40 OD010440). The *C. elegans* MYb minimal native microbiome was a gift from Hinrich Schulenburg. *Salmonella enterica* LT2 was obtained from ATCC (ATCC 700720). *Staphylococcus aureus* MSSA Newman was a gift from Chris LaRock (Emory University).

### Worm colonization and CFU counting

Nematodes were grown, maintained, and manipulated using standard techniques (Stiernagle 2006). Briefly, breeding stocks were maintained on NGM plates + OP50 at 25°C and synchronized using a standard bleach/NaOH protocol where eggs were allowed to hatch out in M9 worm buffer overnight (∼16h) with shaking (200 RPM) at 25°C. Starved L1 larvae were transferred to 10cm NGM plates containing lawns of *E. coli pos-1* RNAi and incubated at 25°C for 3 days to produce reproductively sterile adults. Worms were then transferred to liquid S-medium + 200 µg/ml gentamicin + 50 µg/ml chloramphenicol + 2X heat-killed OP50 (to trigger feeding) for 24-48 hours, resulting in largely germ-free adults. Adult worms were washed via sucrose floatation before colonization.

For colonization, *S. aureus* and *S. enterica* were grown from glycerol stocks to stationary phase overnight in 1 mL LB media at 37°C with shaking at 200 RPM. 10 cm NGM agar plates were seeded with 100 µL of culture, which was distributed with plating beads to cover the surface and incubated 24 hours at 25°C to form lawns. Adult worms prepared as described above were aliquoted onto plates in 25-50 µL drops of buffer (M9 worm buffer + 0.1% Triton X-100 to prevent sticking; ∼500-1000 worms/plate) and incubated at 25°C in the dark for 24 hours (*S. aureus*) or 48 hours (*S. enterica*) to allow colonization. Colonized worms were prepared for mechanical disruption as previously described, then sorted individually or in batches into wells of a deep 2 mL 96-well plate (Axygen) using a BioSorter large object sorter fitted with a 250 µM FOCA (Union Biometrica). Sorted worms were disrupted in 96-well plate format as previously described (Taylor and Vega 2021; Taylor *et al*. 2022a).

Minimal community colonization data (Taylor and Vega 2021) and colonization data for *Microbacterium oxydans* (Taylor *et al*. 2022b) were previously published, and methods are available from the indicated manuscripts.

### Data analysis and simulations

All data analysis and simulations were performed in R 4.2.3 using structures from base R (R Core Team 2022) and tidyverse (Wickham *et al*. 2019). Differences between groups were assessed using functions *t.test()* and *wilcox.test()* from *stats* to perform two-sided testing against a null hypothesis of no difference. Calculation of higher moments of data used functions *skewness()* and *kurtosis()* from package *e1071* (Meyer *et al*. 2021). Gaussian mixture models were fitted to log_10_-transformed data using package *mclust* (Scrucca *et al*. 2016). Plotting was carried out using functionality from packages *ggplot2* (Wickham 2016 p. 2), *cowplot*, (Wilke 2020) and *ggpubr* (Kassambara 2020).

Bootstrapping was done on raw colony count data. For each worm in a data set, the data are a number of counted colonies and a ten-fold serial dilution factor (e.g. 25 colonies counted at the second ten-fold dilution of the original sample). Within each run of the bootstrap, the original data set was resampled with replacement to produce a user-specified number of batch-based data points.

## Data availability

Strains are available upon request. All code and data used in this manuscript are available on GitHub (https://github.com/veganm/WormCFUHeterogeneity2022).

