## Supplemental File S1 for "Sample pooling inflates error rates in between-sample comparisons: an empirical investigation of the statistical properties of count-based data"

**Supplementary Information File 1**

**Multimodality of CFU/worm data**

One qualitative difference that we observe between samples in bacterial load data is the position and/or shape of modes or high-density regions. These data frequently show more than one mode; two modes are the most common. For example, fitting a Gaussian mixture model to the full set (three replicates) of S*almonella enterica* log_10_(CFU/worm) data best supports a two-mode model (**Figure S1**), with (mean_1_, var_1_)=(3.22, 0.19) and (mean_2_, var_2_)=(4.66, 0.19), containing 39% and 61% of the mass respectively.

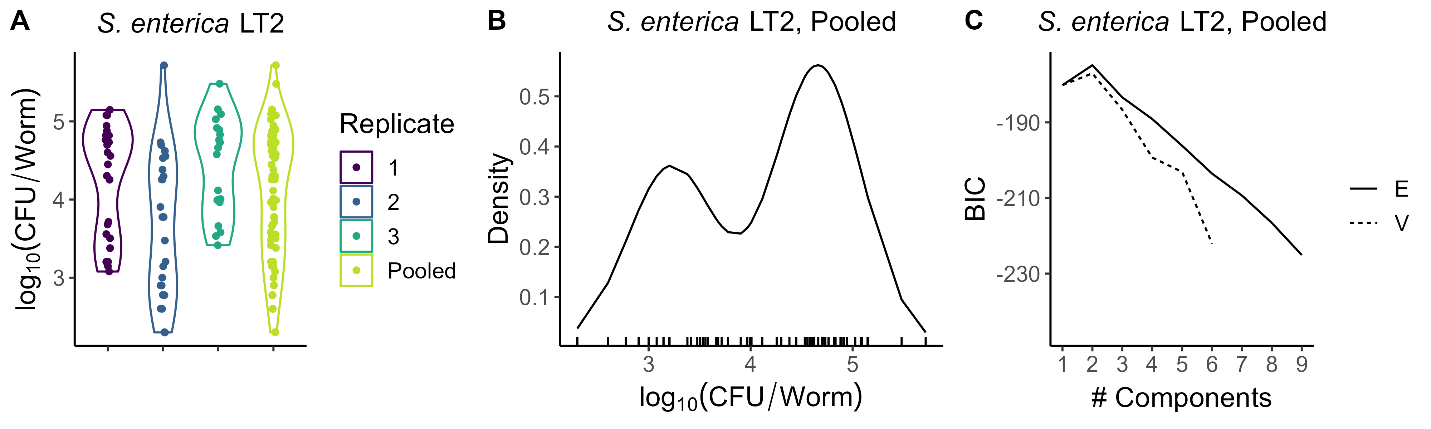

*Figure S1. More than one mode in colonization of N2 adults by S. enterica LT2. A) Three runs of single-worm CFU data (with zeros removed). Runs 1 and 2 were separate experiments two weeks apart; run 3 data were taken a year later by a different operator. (B) Density curve of the two-mode Gaussian mixture model with unequal variances fit to log_10_-transformed pooled data set including all three replicates, with individual data points in x-axis rugplot. (C) BIC plot showing support for GMM fits to pooled data with 1-9 components, assuming equal (“E”) or unequal (“V”) variance.*

It is easy to demonstrate that multimodality by itself is insufficient to generate false positives. In simulations, we generated log-scale data from a two-component Gaussian mixture model using the parameters obtained from *Salmonella* data and created simulated “batches” of 1-50 individual worms. As a sanity check, when two sets of simulated data were created from the same parameter set, these simulations produced the expected false positive rate of ~0.05 regardless of batching (100 runs, 50 points per run per batch size; **Figure S2, Table S1A**).

Allowing run-to-run variation produced the expected batch-induced inflation in statistically significant comparisons between "replicates". For example, fitting replicate 1 data to a two-component GMM by itself generated a model with two modes: (mean_1_, var_1_)=(3.4, 0.06) and (mean2, var2)=(4.8, 0.06), containing 34.6% and 65.5% of the mass respectively. For replicate 3, the model again supported two modes, with (mean_1_, var_1_)=(3.8, 0.06) and (mean_2_, var_2_)=(4.9, 0.06), containing 42.1% and 57.9% of the mass respectively. As the positions of the modes were similar to one another and to the fit to the full data set, for simplicity we used those GMM parameters for simulation, while allowing the proportion of the population within each mode to vary as observed in these data (~10% difference, **Figure S2**). In these simulations, batching amplified differences between the run 1 and run 2 data and increased the likelihood of rejecting the null hypothesis of no difference across runs (**Table S1B**). With a smaller difference in proportions (5%, **Figure S2C**), similar results were obtained, albeit less pronounced (**Table S1C**). Note that in these simulations the results of unbatched data comparisons can be thought of as analogous to power (how often testing detects the difference between groups).

| **(A) Pooled data parameterization** | | | | | |
| --- | --- | --- | --- | --- | --- |
| Batch size | 1 | 5 | 10 | 20 | 50 |
| t-test | 0.05 | 0.05 | 0.08 | 0.09 | 0.05 |
| Wilcoxon | 0.03 | 0.03 | 0.08 | 0.08 | 0.06 |
| **(B) Pooled data parameterization, 10% difference in mode proportions** | | | | | |
| Batch size | 1 | 5 | 10 | 20 | 50 |
| t-test | 0.12 | 0.32 | 0.51 | 0.64 | 0.94 |
| Wilcoxon | 0.13 | 0.28 | 0.49 | 0.64 | 0.94 |
| **(C) Pooled data parameterization, 5% difference in mode proportions** | | | | | |
| Batch size | 1 | 5 | 10 | 20 | 50 |
| t-test | 0.09 | 0.11 | 0.15 | 0.27 | 0.36 |
| Wilcoxon | 0.06 | 0.10 | 0.13 | 0.22 | 0.33 |

*Table S1. Fraction of tests with p>0.05 in comparisons of simulated data (n=50 points per condition, 100 simulation runs) from a Gaussian mixture model with two modes. Parameters were obtained by fitting a two-component GMM to the pooled log_10_CFU Salmonella data in* ***Figure S1*** *((mean_1_, var_1_)=(3.22, 0.19) and (mean_2_, var_2_)=(4.66, 0.19), containing 39% and 61% of the mass respectively). In all simulations, proportion of data in each mode is stochastic (membership of each individual simulated worm is drawn from a binomial distribution, using the probabilities indicated). (A) Random variation only, using parameter estimates from data. The same parameters are used for purple and orange data “replicates”; differences between “replicates” are from the simulation. (B) Parameters for purple/orange data are identical (same as in (A)) but proportions in modes differ by 10% between replicates (purple, low mode contains 39% of the mass; orange, 29%). (C) Parameters are as before; 5% difference in proportions in modes. One example simulation for each scenario is shown in* ***Figure S2****.*

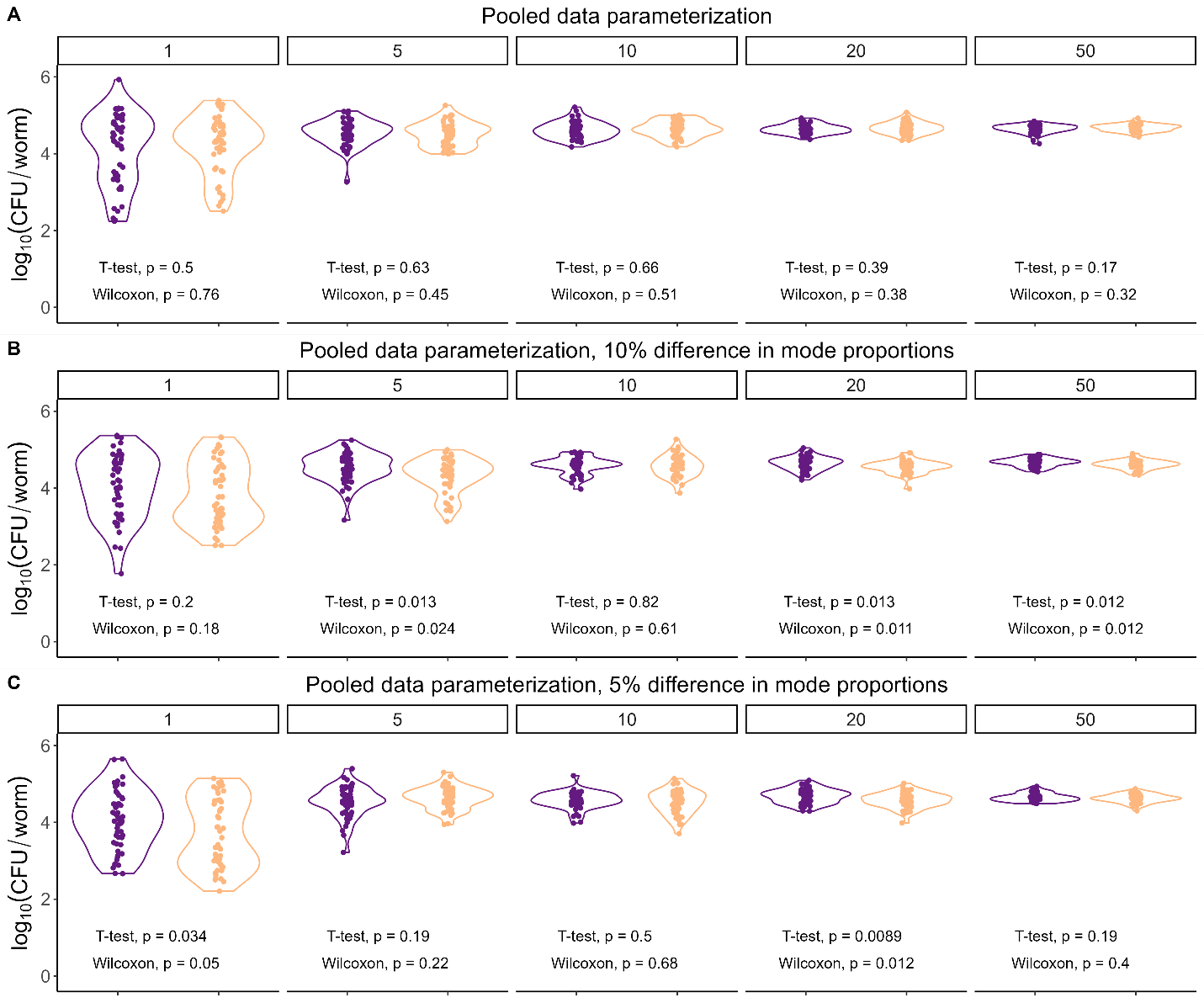

*Figure S2. One example of simulated data (n=50 points in each data set, two replicates) from a Gaussian mixture model with two modes. Labels above each panel indicate the number of simulated individuals per batch. Parameters were obtained by fitting a two-component GMM to the pooled log_10_CFU Salmonella data in* ***Figure S1*** *((mean_1_, var_1_)=(3.22, 0.19) and (mean_2_, var_2_)=(4.66, 0.19), containing 39% and 61% of the mass respectively). In all simulations, proportion of data in each mode is stochastic (membership of each individual simulated worm is drawn from a binomial with the probabilities indicated). (A) Random variation only, using parameter estimates from data. The same parameters are used for purple and orange data “replicates”; differences between “replicates” are from the simulation. (B) Parameters for purple/orange data are identical (same as in (A)) but proportions in modes differ by 10% between replicates (purple, low mode contains 39% of the mass; orange, 49%). (C) Parameters are as before; 5% difference in proportions in modes. Results are shown for comparisons between replicates; statistics for 100 runs of the simulation are shown in* ***Table S1****.*

**Effects of variation and skew on false-positive rates**

As real CFU/worm data tend to be right-skewed and highly variable, often covering two or more orders of magnitude, we performed simulations to determine whether generation of false positives is related to these higher moments of the data. The beta distribution is a flexible distribution with desirable properties for these simulations. The distribution is on the range [0,1], and it is easy to move data into a desired range using a constant multiplier (here, 10^5^ to bring these data into the range of worm CFU counts). As the mean E[x] for a beta distribution is a simple function of the two parameters $E\left[ x \right]=\frac{\alpha}{\alpha+\beta}=\frac{1}{1+\beta/\alpha}$, distributions with the same mean but different higher moments can be produced by holding the ratio β/α constant (**Figure S3**).

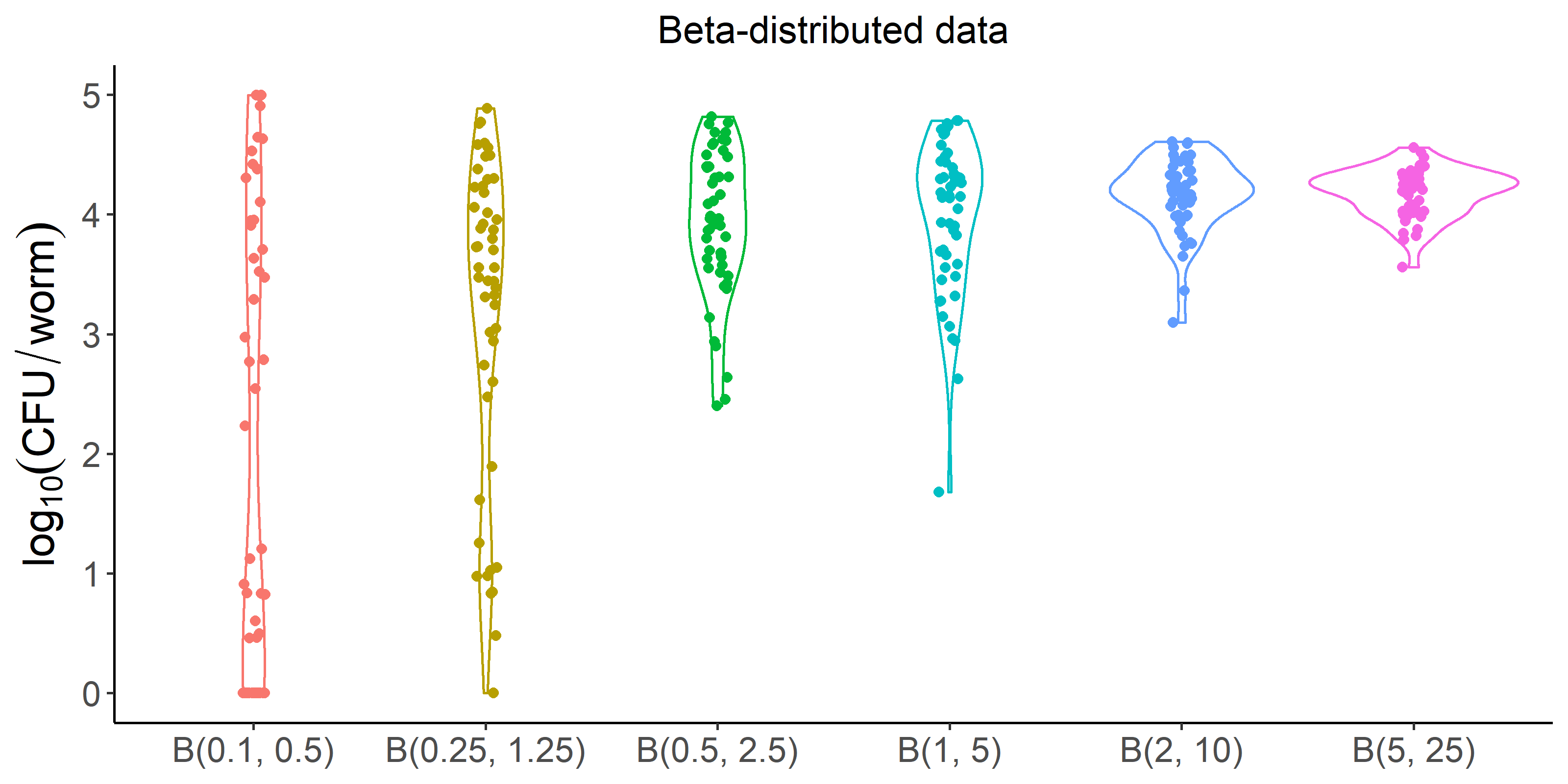

*Figure S3. Simulated data based on the beta distribution. In these simulations, we start with Beta(0.1, 0.5) and alter the distribution of the data by multiplying both α and β by a constant factor (2.5-50). Simulated data (n=50 per distribution) are multiplied by a constant 10^5^ to produce values expected for CFU/worm. Note that all distributions have the same pre-transformation mean of 1/6 and post-transformation mean of ~16,667.*

Over the parameter range used in **Figure S3**, the distribution goes from bimodal with high variance and skew (α<1, β<1) to fairly condensed and nearly symmetric (**Table S2**). The expectation in all cases was 10^5^*(1/6) = 16,667, and the averages of all sets of simulated data (n=50 per distribution) were close to this value. All these distributions were right skewed (median < mean), since α < β. However, the distance between the median and the average became smaller as α and β increased (decreasing skew), as did the variance. In testing for differences across these data sets, the nonparametric Wilcoxon test (which compares the rank sum of values from both distributions) could in many cases tell these data apart, but the two-sample t-test (which compares the means) very reasonably could not (**Table S3**).

|  | **β(0.1, 0.5)** | **β(0.25, 1.25)** | **β(0.5, 2.5)** | **β(1,5)** | **β(2, 10)** | **β(5, 25)** |
| --- | --- | --- | --- | --- | --- | --- |
| **Average** | 12,521 | 11752 | 17,262 | 19,020 | 17,753 | 16,716 |
| **Median** | 14 | 3579 | 9,198 | 14.264 | 15,853 | 16,984 |

*Table S2. Summary statistics for beta-distribution derived simulated data.*

|  | **β(0.1, 0.5)** | **β(0.25, 1.25)** | **β(0.5, 2.5)** | **β(1,5)** | **β(2, 10)** | **β(5, 25)** |
| --- | --- | --- | --- | --- | --- | --- |
| **β(0.1, 0.5)** |  | 3.20E-03 | 7.72E-06 | 3.75E-06 | 3.13E-07 | 3.25E-07 |
| **β(0.25, 1.25)** | 0.86 |  | 0.01 | 2.39E-03 | 3.58E-05 | 1.83E-05 |
| **β(0.5, 2.5)** | 0.30 | 0.12 |  | 0.63 | 0.08 | 0.06 |
| **β(1,5)** | 0.16 | 0.04 | 0.62 |  | 0.39 | 0.38 |
| **β(2, 10)** | 0.20 | 0.04 | 0.86 | 0.67 |  | 0.96 |
| **β(5, 25)** | 0.29 | 0.07 | 0.84 | 0.40 | 0.55 |  |

*Table S3. Results of pairwise comparisons between the sets of simulated data shown in Figure. Top diagonal of the matrix contains Wilcoxon test p-values; bottom diagonal contains t-test p-values. No correction for multiple comparisons was performed, as this does not alter the qualitative differences between tests. Color of each cell indicates distance from p=0.05 for reference only.*

This presents an interesting situation. For the beta-distributed "individual data", the parametric test ignores the effects of skew in deciding (correctly) that these "populations" have the same mean, whereas the nonparametric test treats the distribution as information and concludes (still correctly) that many of these "populations" are different from one another. Both statements are true; as these tests ask different questions, they arrive at different conclusions. As batching effectively creates an arithmetic mean across individuals within a batch, this feature of the tests will be useful for understanding why batching can alter comparisons.

For the purposes of illustration, consider a pair of distributions that provide a roughly similar range of values as in the *Salmonella* single-worm data: Beta(0.5, 2.5) and Beta(1,5) (**Tables S2-S3**). Recall that these distributions do have the same means, and that in the "individual data" comparison in **Table S3**, neither pairwise test rejected the null hypothesis of no difference for this pair of distributions. As a sanity check, creating two sets of simulated data representing the same distribution and using batch sizes of (1, 5, 10, 20, 50) resulted in false-positive rates that were close to the expected α=0.05 (over 1000 runs, **Table S4**). Next we created sets of simulated data from pairs of different distributions (e.g. Beta(0.5, 2.5) vs Beta(1, 5)). The nonparametric test was sometimes able to reject the null hypothesis, but this was lost as batch size increased; meanwhile, the parametric test correctly failed to discern a difference between sample means regardless of whether batching was used. Drawing from two distributions farther apart in parameter space (Beta(0.5, 2.5) vs Beta(2, 10)), the nonparametric test rejected the null somewhat more often (larger signal = higher power), but this was once again lost when comparing batched samples (**Table S4**).

| **Beta(0.5, 2.5) vs Beta(0.5, 2.5)** | | | | | |
| --- | --- | --- | --- | --- | --- |
| **Batch** | **1** | **5** | **10** | **20** | **50** |
| **t-test** | 0.053 | 0.055 | 0.047 | 0.051 | 0.05 |
| **Wilcoxon** | 0.045 | 0.052 | 0.058 | 0.047 | 0.044 |
| **Beta(1, 5) vs Beta(1, 5)** | | | | | |
| **Batch** | **1** | **5** | **10** | **20** | **50** |
| **t-test** | 0.036 | 0.049 | 0.057 | 0.037 | 0.044 |
| **Wilcoxon** | 0.031 | 0.048 | 0.058 | 0.042 | 0.045 |
| **Beta(0.5, 2.5) vs Beta(1, 5)** | | | | | |
| **Batch** | **1** | **5** | **10** | **20** | **50** |
| **t-test** | 0.051 | 0.063 | 0.051 | 0.064 | 0.044 |
| **Wilcoxon** | 0.113 | 0.050 | 0.046 | 0.062 | 0.043 |
| **Beta(0.5, 2.5) vs Beta(2, 10)** | | | | | |
| **Batch** | **1** | **5** | **10** | **20** | **50** |
| **t-test** | 0.052 | 0.088 | 0.072 | 0.059 | 0.052 |
| **Wilcoxon** | 0.211 | 0.061 | 0.059 | 0.060 | 0.054 |

*Table S4. Pairwise comparisons of randomly-generated data at different levels of batching (n=1, 5, 10, 20, or 50 individuals/batch), drawn from the indicated pairs of distributions. Values given are the fraction of 1000 runs where p≤0.05 for each test.*

These simulations provide an illustration of the fact that batching across individuals behaves mathematically as an average, with all the properties that entails. If two sets of data are from populations with the same mean, any comparison of those data that relies on the properties of the population mean (here, t-tests or comparisons of batches) rather than the overall distribution should see those populations as indistinguishable. In this case, the result was a false negative for the actual hypothesis of interest - for batched data drawn from two populations with very different distributions but a shared mean, these tests consistently (and reasonably) failed to reject the null hypothesis of no difference when comparing the data averages.

*Effects of run-to-run variation*

Next we sought to determine how typical run-to-run variation would affect comparisons of batch-averaged data. These simulations used Beta(1,5) as a baseline and additionally assumed some run-to-run variation in the parameters from which the data are generated. Specifically, new parameters were generated for each data set *i*=(A, B) in each run as $\left( \alpha_{i},\beta_{i} \right)=\left( \alpha\left( 1+U\left( -0.1, 0.1 \right) \right), \beta\left( 1+U\left( -0.1, 0.1 \right) \right) \right)$. Allowing run-to-run variation in parameters produced inflated false-positive rates when comparing biologically averaged “batch” data (see main text and **Table 1**). The between-group distances for moments of the single-"worm" data in these simulations (**Figure S4)** were consistent with those observed in real data (**Figure 2**). Higher moments of the simulated data affected the distribution of between-group distances, as can be seen by comparing data simulated from Beta(1,5) (var 0.02 and skewness 1.18) vs. those simulated from Beta(5,5) (var 0.02 and skewness 0) (**Figure S5**).

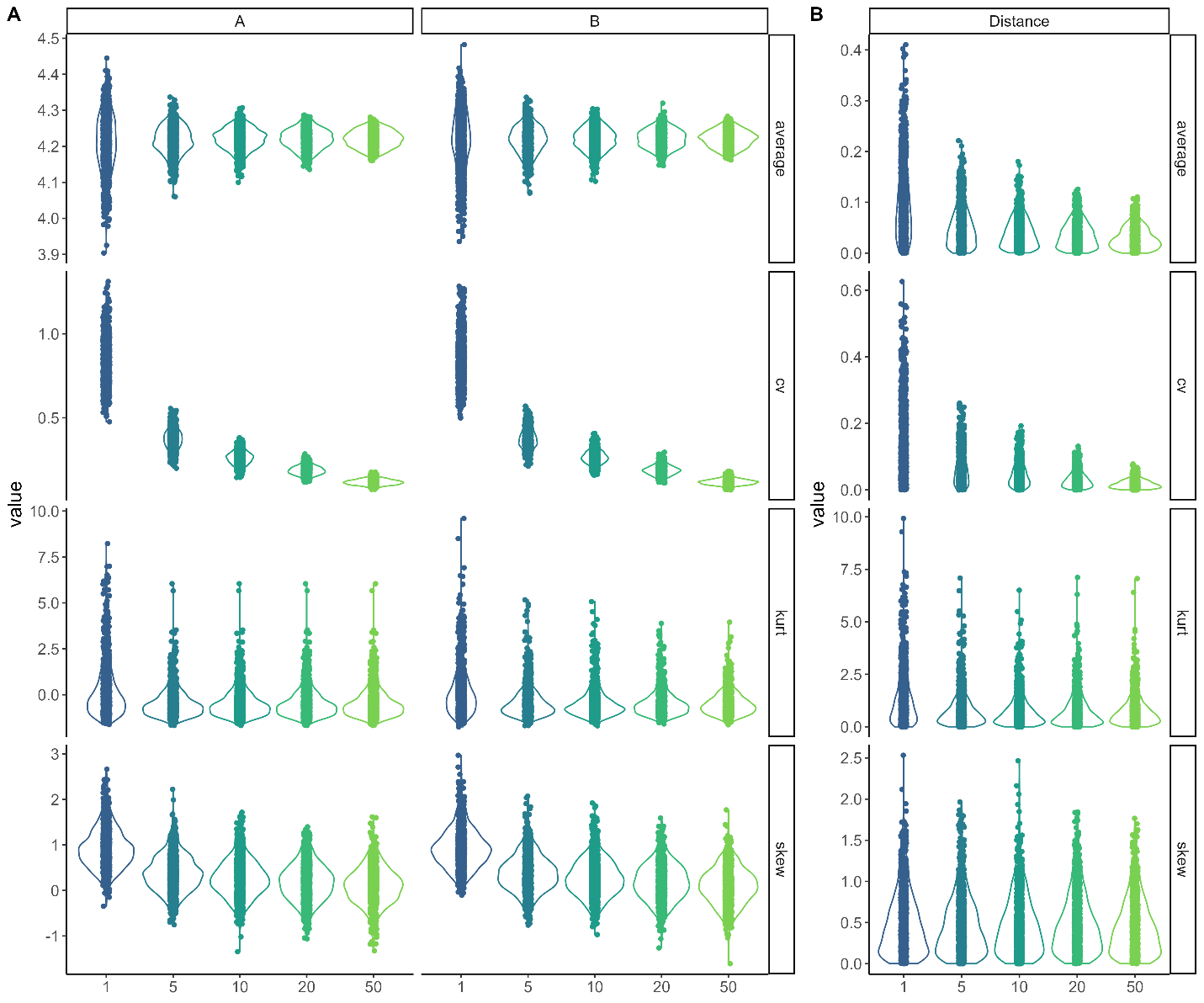

*Figure S4. Trends in moments in simulated data (1000 runs, max 10^5^ CFU/worm), where two random samples (A and B, 24 data points for each sample at each batch size) are generated from distributions Beta(α_A_,β_A_) and Beta(α_B_,β_B_) respectively within each run, with shared baseline distribution Beta(1,5) and parameter deviances drawn from U(-0.1, 0.1). Batch size is on the x-axis in all plots. (A) Summary statistics over n=1000 simulations for simulated data sets A and B. (B) Distances between moments of the data sets A and B within each run of the simulation.*

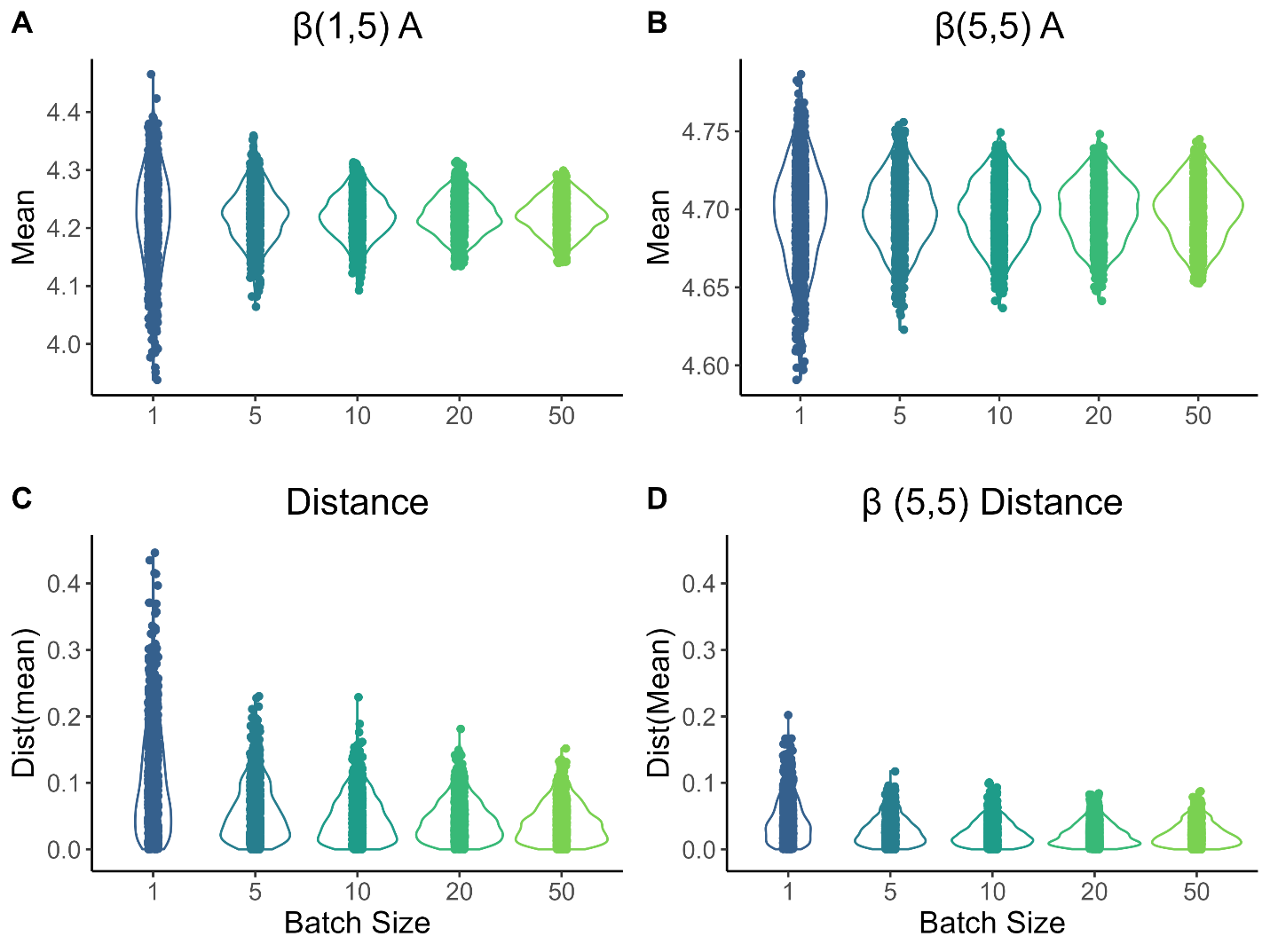

*Figure S5. Distribution of population means (A,B) and distances between means (C,D) for simulated replicates from the same underlying distribution. (A, C) Beta(1,5); (B, D) Beta(5,5). Data were simulated using 24 data points at each batch size per run, 1000 runs, maximum value 10^5^ CFU/worm.*

*Effects of replication*

We next modified our simulations to include replicate runs in each data set in each run of the simulation (3, 10, or 25 replicates), where randomized parameters were re-drawn each “run”, and to include limitations on both the number of digests performed and the total number of individuals available. In these simulations, the total number of simulated “worms” was set as a constant, and the indicated number of worms were divided up into batches of indicated size. This meant that the total number of data points decreased with batch size (e.g. 100 total worms could be divided into 20 batches of 5 worms each but only two batches of 50).

We show the effects of run-to-run variation in parameters in experiments of various sizes: (1) “small” (100 total worms, maximum of 12 data points per run regardless of batch size), (2) “medium” (200 total worms, maximum of 24 data points per run), or (3) “large” (500 total worms, maximum of 48 data points per run) (**Table S5**). As before, false-positive results are inflated due to batching. While we observe decreased false positives in small experiments and at high batch sizes, this is due to loss of power rather than to mitigation of the effects of batching (in the “small” simulations with 100 worms, only two batches of 50 worms can be created!). If we instead allow n=24 data points regardless of batch size, we restore the monotonic relationship between batch size and false positive rate (**Table S5**). Replication improves the within-experiment distances between samples (**Figures S6, S7**), but the effect is small.

As before, symmetry increases false positives. Allowing run-to-run variation when drawing values from the symmetric distribution Beta(5,5) instead of Beta(1,5) (similar variance) readily creates apparent differences between groups, and as before, batching further exacerbates these differences. Drawing instead from the intermediate-skew distribution Beta(3,7) (variance 0.019, skewness 0.48) is of some help (**Table S5**). Overall, although replication is helpful in measuring run-to-run variation in real experiments and is expected to increase power to detect true differences, we find that replication does not materially affect false-positive rate inflation due to batching.

|  |  | **Batch Size** | | | | |
| --- | --- | --- | --- | --- | --- | --- |
| **Distribution** | **Condition** | **1** | **5** | **10** | **20** | **50** |
| B(1,5) | small | 0.06 | 0.08 | 0.11 | 0.09 | 0.07 |
| B(1,5) | medium | 0.06 | 0.13 | 0.17 | 0.17 | 0.15 |
| B(1,5) | large | 0.09 | 0.21 | 0.35 | 0.33 | 0.30 |
| B(1,5) | mediumx3 | 0.07 | 0.15 | 0.21 | 0.30 | 0.46 |
| B(1,5) | mediumx10 | 0.06 | 0.15 | 0.18 | 0.29 | 0.45 |
| B(1,5) | mediumx25 | 0.08 | 0.14 | 0.22 | 0.31 | 0.44 |
| B(1,5) | medium,n=24 | 0.07 | 0.13 | 0.22 | 0.33 | 0.49 |
| B(5,5) | small | 0.09 | 0.18 | 0.21 | 0.19 | 0.11 |
| B(5,5) | medium | 0.10 | 0.25 | 0.33 | 0.29 | 0.23 |
| B(5,5) | large | 0.13 | 0.36 | 0.49 | 0.46 | 0.41 |
| B(5,5) | medium,n=24 | 0.10 | 0.29 | 0.38 | 0.53 | 0.66 |
| B(5,5) | mediumx3 | 0.10 | 0.25 | 0.35 | 0.46 | 0.57 |
| B(5,5) | mediumx10 | 0.10 | 0.27 | 0.38 | 0.46 | 0.60 |
| B(5,5) | mediumx25 | 0.09 | 0.25 | 0.35 | 0.45 | 0.58 |
| B(3,7) | small | 0.06 | 0.14 | 0.17 | 0.14 | 0.07 |
| B(3,7) | medium | 0.09 | 0.24 | 0.32 | 0.28 | 0.21 |
| B(3,7) | large | 0.14 | 0.36 | 0.48 | 0.47 | 0.38 |
| B(3,7) | medium,n=24 | 0.10 | 0.25 | 0.36 | 0.52 | 0.65 |

*Table S5. Including features of experimental realism fails to resolve false-positive rate inflation. Data shown are the fraction of runs where Mann-Whitney U test p<0.05 (over 1000 runs, in order of batch size). For indicated simulations, the total number of worms is constrained such that the number of data points decreases with batch size: (1) “small” (100 total worms, maximum of 12 data points per run regardless of batch size), (2) “medium” (200 total worms, maximum of 24 data points per run), or (3) “large” (500 total worms, maximum of 48 data points per run). For simulations with multiple replicates (x3, x10, or x25 “runs” of sampling), or for simulations with a single replicate run when indicated (n=24), all batch sizes are represented by 24 data points per run.*

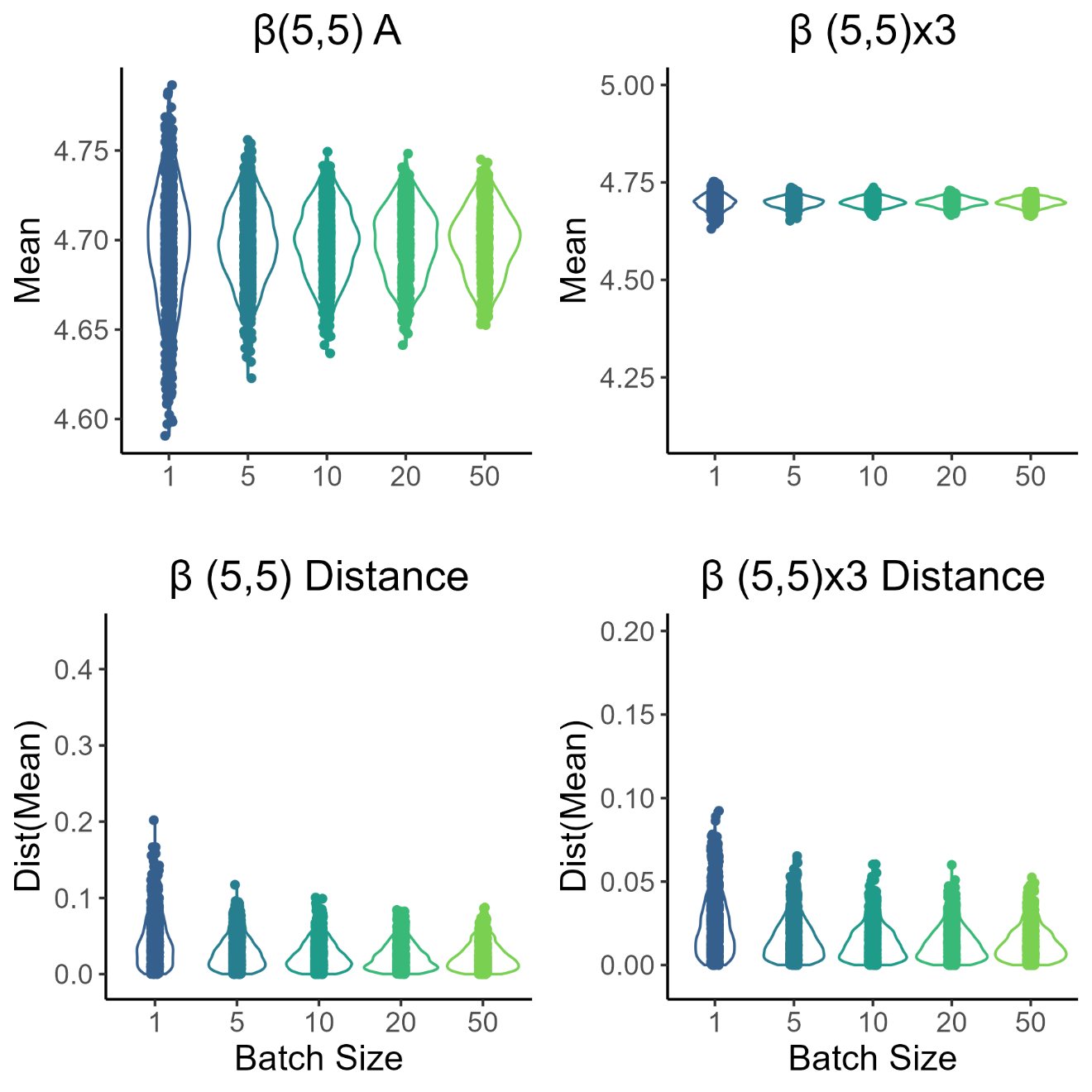

*Figure S6. Replication slightly improves within-experiment distances between samples drawn from the same base distribution with error. Average values and normalized distances of between-sample means (10,000 runs, n=24 data points per run at all batch sizes) for Beta(5,5) single-run vs three-replicate simulations. Replication somewhat tightens the distribution of the means within samples, and the distribution of between-sample distances is weakly affected by batching.*

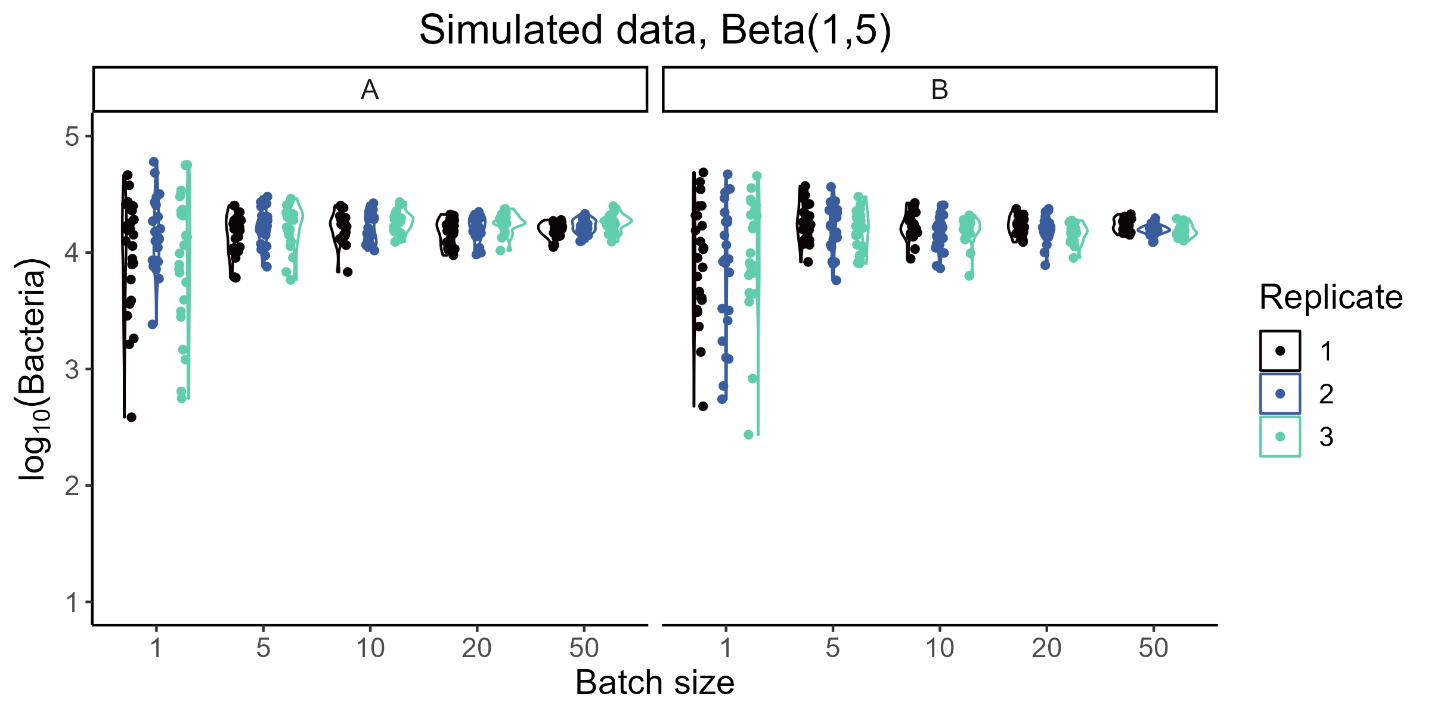

*Figure S7. Data from one example simulation with n=24 data points at each batch size, where all data were simulated from the underlying distribution Beta(1,5) with deviation on both parameters drawn from U[-0.1, 0.1]. New parameters were generated for each “condition” (A, B) on each of three “replicates”.*
